# MRI-Guided Focused Ultrasound Blood-Brain Barrier Opening Increases Drug Delivery and Efficacy in a Diffuse Midline Glioma Mouse Model

**DOI:** 10.1101/2023.04.05.534448

**Authors:** Payton Martinez, Genna Nault, Jenna Steiner, Michael F. Wempe, Angela Pierce, Breaunna Brunt, Mathew Slade, Andrew Mongin, Jane Song, Kang-Ho Song, Nicholas Ellens, Natalie Serkova, Adam Green, Mark Borden

## Abstract

Diffuse intrinsic pontine glioma (DIPG) is the most common and deadliest pediatric brainstem tumor and is difficult to treat with chemotherapy in part due to the blood-brain barrier (BBB). Focused ultrasound (FUS) and microbubbles (MBs) have been shown to cause BBB disruption (BBBD), allowing larger chemotherapeutics to enter the parenchyma. Panobinostat is an example of a promising *in vitro* agent in DIPG with poor clinical efficacy due to low BBB penetrance. In this study, we hypothesized that using FUS to disrupt the BBB allows higher concentrations of panobinostat to accumulate in the tumor, providing a therapeutic effect. Mice were orthotopically injected with a patient-derived DMG cell line, BT-245. MRI was used to guide FUS/MB (1.5 MHz, 0.615 MPa PNP, 1 Hz PRF, 10 ms PL, 3 min treatment time) / (25 µL/kg, IV) targeting to the tumor location. In animals receiving panobinostat (10 mg/kg, IP) in combination with FUS/MB, a 3-fold increase in tumor panobinostat concentration was observed, with only insignificant increase of the drug in the forebrain. In mice receiving three weekly treatments, the combination of panobinostat and FUS/MB led to a 71% reduction of tumor volumes by MRI (*p* = 0.01). Furthermore, FUS/MB improved the mean survival from 21 to 31 days (*p* < 0.0001). Our study demonstrates that FUS-mediated BBBD can increase the delivery of panobinostat to an orthotopic DMG tumor, providing a strong therapeutic effect and increased survival.

**One Sentence Summary:** FUS and microbubbles can increase the delivery of panobinostat to a patient-derived xenograft (PDX) orthotopic DMG tumor, providing a strong therapeutic effect and increased survival.

## INTRODUCTION

In 1926, Wilfred Harris first described diffuse intrinsic pontine glioma (DIPG) affecting the pontine area of the brainstem. To this day, DIPG remains one of the most difficult brain tumors to treat^1^. Imaging is crucial for establishing DIPG diagnosis, with magnetic resonance imaging (MRI) considered as the gold standard^2^. Further classification is made through genetic sequencing of biopsies taken from patients; it has been shown that over 80% of DIPG share a similar somatic gain-of-function mutation, where a missense substitution is found for lysine 27 to methionine in either histone variant HIST1H3B (H3.1) or H3F3A (H3.3)^3–5^. The World Health Organization (WHO) recently classified DIPGs and other midline tumors with this mutation as “H3 K27M-altered diffuse midline gliomas (DMG)“^6^. With this information, many cytotoxic^7,8^ and targeted^9,10^ drugs have been the subject of DMG clinical trials. Unfortunately none of these drugs have worked in clinical studies owing in part to the impermeability of the tumor blood brain barrier (BBB)^11,12^.

One major obstacle to the efficacy of current treatments is the low accumulation of chemotherapeutic drugs in the tumor region. The pons, where most DMG are located, has an intact BBB that prevents most chemotherapies from entering the parenchyma. Moreover, the pons possesses a more unyielding BBB, further complicating the penetration of pharmaceuticals^13^. An impermeable blood brain tumor barrier (BBTB) also limits the tumor margin visibility on T1-weighted contrast enhanced MRI scans^14^. A study by McCully et al. showed that the BBTB is not homogeneous, as there was a difference in temozolomide penetration between the brainstem and pons region relative to the cortex^13^. While there are a select few chemotherapeutic agents, including gemcitabine, that have shown the ability to penetrate the BBTB in a human pontine DMG^15^, unfortunately these agents have not improved outcomes. Therefore, delivery methods are needed to enhance drug efficacy in these tumors.

One drug delivery technique that has been used to address this issue is convection enhanced delivery (CED). CED involves the insertion of a catheter through the skull into the pons region in this case, from which drugs are pumped using positive pressure and an engineered catheter tip to enhance convective mass transport into the surrounding tissue^16,17^. This method has shown higher therapeutic penetrance at the target site with less systemic toxicity^16,17^. However, CED can present complications, especially in the treatment of solid tumors. Some tumors are highly vascularized, develop increased interstitial pressure, and are less susceptible to the pressure-driven approach^18^. Alternatively, quickly growing tumors can develop areas of necrosis, rendering CED less effective as drugs can pool in this area preventing delivery to the faster-growing cells on the periphery^19^. The invasiveness of this technique is also a concern, as it requires stereotactic surgery to pierce through the scalp, skull, dura and intervening healthy brain tissue to place the catheter(s) in the tumor. Furthermore, clinical trials have shown the potential for an increase in neurological damage at higher CED flow rates^20^.

Focused ultrasound complemented with microbubbles (FUS/MB) has materialized as a non-invasive technique to momentarily disrupt the BBB in a targeted and reversible fashion^21–24^. MBs utilized for this technique were originally developed and clinically approved as ultrasound contrast agents. They are typically 1-10 µm diameter, with a shell (e.g., lipid or protein) that encapsulates a high-molecular-weight internal gas (e.g., perfluorocarbon)^25–27^. The physical mechanism is that the circulating, gas-filled microbubbles expand and contract (cavitate) under the exposure of ultrasonic pressure waves (0.2-2 MHz) far more than the surrounding fluid and viscoelastic tissue. As a result, the local mechanical forces cause the separation of tight junctions, disrupting the BBB^28–31^. Researchers have classified two types of cavitation based on their acoustic echo: harmonic and inertial. Harmonic cavitation is characterized by relatively small oscillations of the microbubble and produces a frequency response at harmonics of the driving frequency^32,33^. The sub and ultra-harmonic responses are caused by the nonlinear and asymmetric response of the microbubbles as they expand and contract^32^. Inertial cavitation can be described by more violent expansion and contraction and produces a broadband frequency response due to shock waves generated by MB implosions^33^. Both regimes of cavitation produce mechanical forces (e.g., fluid shear stresses, direct contact forces through collision and retraction, acoustic shock waves and micro-jetting) to the endothelium within the focal zone of the FUS, causing sonoporation of the plasma membrane and disruption of the tight junctions^33^.

Passive cavitation detection (PCD) can provide real-time feedback of acoustic activity. Some groups have shown success in using a PCD feedback system that controls the acoustic pressure to maintain a constant cavitation dose^34–36^. FUS-mediated BBB disruption has been shown in clinical trials to be safe, with no significant neuronal damage, apoptosis, ischemia or long-term damage to the vessels^22^. Localized BBB opening can remain for a period of 3 to 24 h, depending on the intensity of the mechanical stresses modulated through acoustic intensity and MB dose^37^. The safety and reversibility of ultrasound-mediated BBB disruption and the small volumetric focal zone attainable makes FUS/MB a good candidate for targeted drug delivery of molecules, particles and cells that cannot otherwise pass the BBB.

We previously identified a strong chemotherapeutic candidate, panobinostat, that has had success against patient-derived xenograft (PDX) DIPG models both in vitro and in vivo^9,38,39^. Although panobinostat is a small molecule drug, it binds to albumin protein once injected and is therefore too large to effectively cross the BBB when injected systemically. Our study uses the novel drug delivery method of FUS/MB guided by high-resolution T2-weighted MRI to deliver panobinostat more effectively to a targeted tumor region and assesses its effectiveness against a patient-derived DIPG tumor model (Fig. 1). The BBBD was assessed by T1-weighted MRI pre-/ post-gadolinium inject, since most of clinically used gadolinium chelates do not penetrate the intact BBB.

**Fig. 1.**
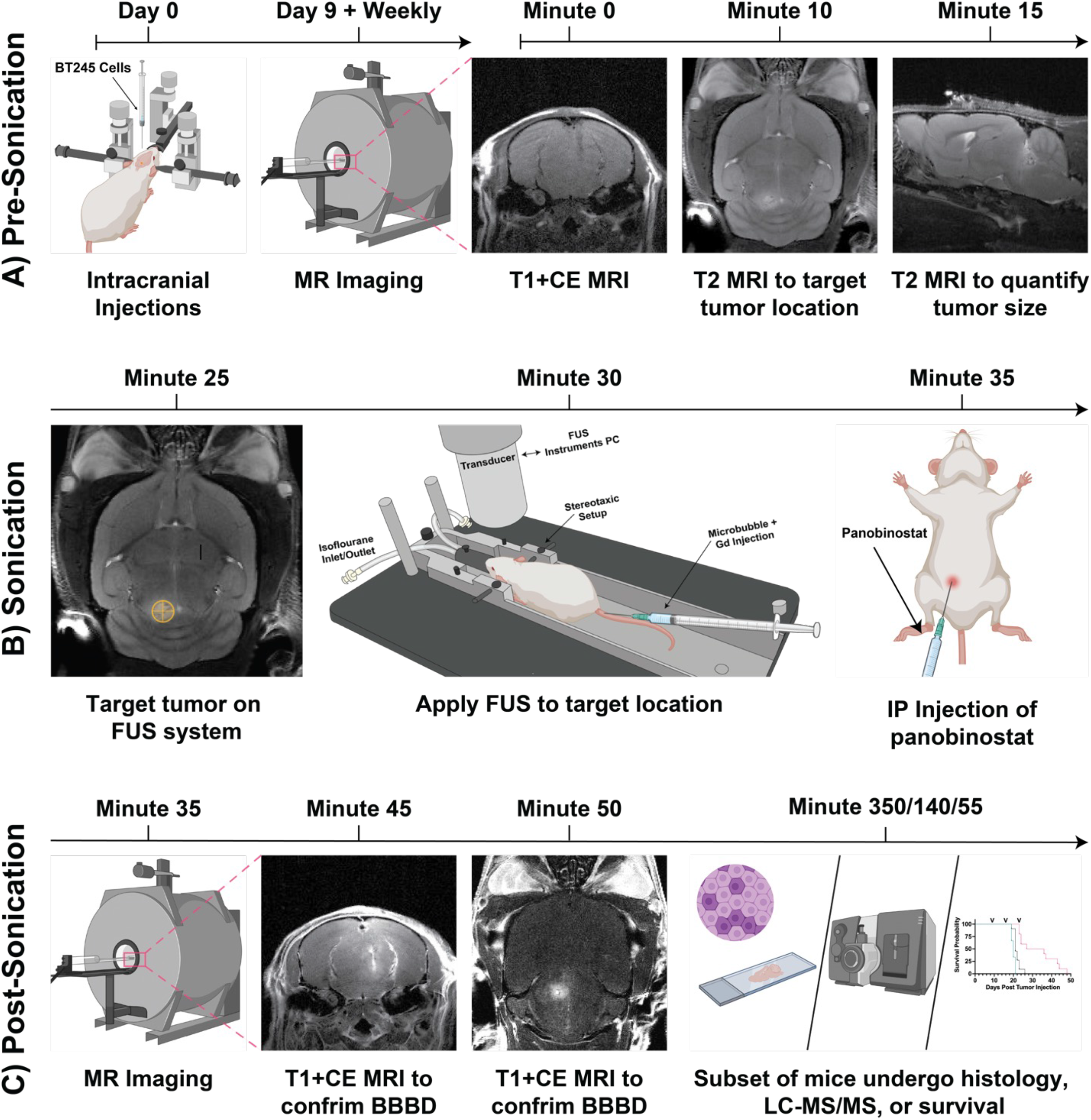
Workflow for our MRI-guided FUS/MB Treatments. (A) Prior to FUS treatment, patient-derived BT245 DIPG cells were injected IC into the pons region. Mice were placed in MRI and axial T1w images, and coronal and sagittal T2w images were acquired. (B) During FUS treatment, coronal T2w MRI images were used to co-register the FUS targeting system. Mice were then moved from MRI on the same bed to the FUS system, and a water tank was placed on the mouse head with ultrasound gel for acoustic coupling between the animal and ultrasound transducer. Microbubbles and gadolinium (Gd) contrast were injected IV, the FUS transducer was stereotactically translated to align the acoustic focus to the pons region of the brain guided by the MRI image, and FUS was applied. Directly after FUS treatment, mice were injected with panobinostat IP. (C) Post FUS, mice were moved back to the MRI to assess BBB opening by T1w MRI scan of Gd extravasation. Mice were then sacrificed and assessed for histology and drug delivery (LC-MS/MS), or housing to continue survival studies. The timeline of study is shown above all images.

## RESULTS

### BT245 xenograft mouse models represent DIPG and DMGs

Among current clinically approved chemotherapeutics, panobinostat was selected for this study as the most effective drug *in vitro* to our patient-derived DIPG cells. To confirm efficacy of panobinostat, BT245 DIPG cells were put through dose escalation trials and MTS assay to determine the half-maximal inhibitory concentrations (IC_50_). The resulting IC_50_ value for panobinostat was 16.1 nM (Fig. 2A). Panobinostat was chosen because it has a low IC_50_ and shows poor BBB penetrance. Most (∼90%) of panobinostat, which associates with albumin via hydrophobic intermolecular forces^40^, is protein bound in blood and prevents its penetration across the BBB or BBTB^41,42^.

**Fig. 2.**
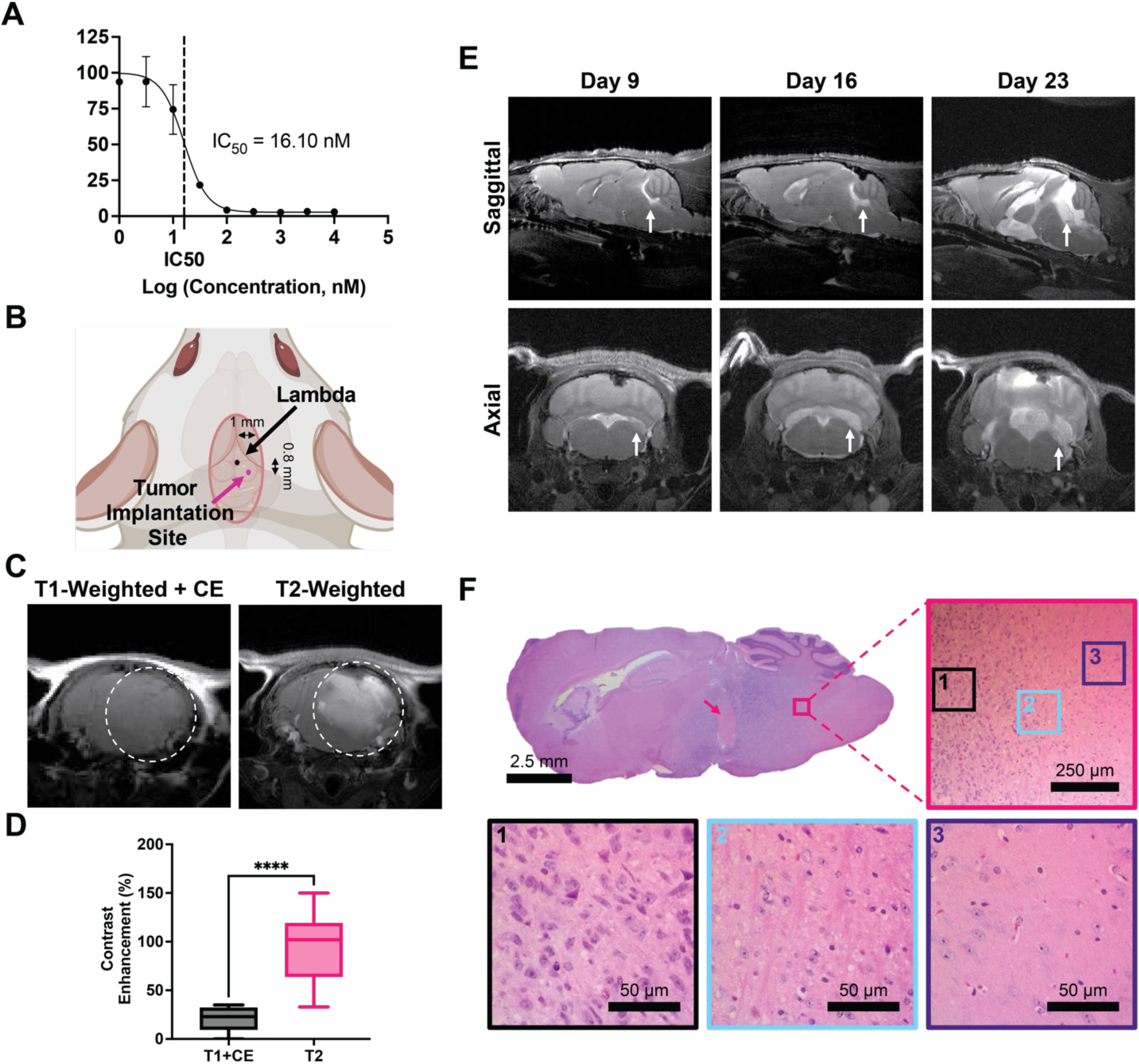
DIPG BT245 PDX model. (A) Cell viability of BT245 cells treated with panobinostat for 72 h assessed by MTS assay. Vertical dotted line represents IC_50_. Values represent the mean ± standard deviation (*n* = 3). (B) Cartoon of tumor implantation site with respect to Lambda skull landmark. (C) T1w contrast-enhanced (CE) MR image (left) prior to FUS/MB showing little tumor enhancement in the T2w MR detectable tumor (right). (D) Quantification of contrast enhancement between the two MRI imaging modes (*n* = 9). (E) T2w MR images at 9, 16 and 23 days post tumor implantation in the control group. Images show intrusive tumors at the pontine region with growth that progresses to the ventricles and adjacent brain structures. (F) Histological analysis of tumor at 3 weeks post tumor implantation. Full view of H&E staining shows whole tumor region, with pink arrow illustrating region of necrosis. Figure is zoomed in to a region on the border of tumor (pink border) and heathy tissue. Also shown are zoomed regions of the tumor (black border), margin (blue border), and healthy tissue (purple border). Abbreviations: MR = Magnetic Resonance; T1w = T1-weighted; T2w = T2-weighted; H&E = Hematoxylin and Eosin. IC_50_ = half maximal inhibitory concentration.

Our PDX orthographic mouse model was established by implanting 2 ×10^5^ luciferase-expressing BT245 cells into the pons of nude athymic mice (Fig. 2B). This cell line has been used extensively in murine models^15,43–45^. Tumor progression was monitored using bioluminescent imaging (BLI) for the first week (Fig. S1). Thereafter, MRI was used to monitor tumor volumes and to guide and validate FUS treatments. One important component of this murine model is the lack of contrast enhancement on T1-weighted (T1w) MR images owing to poor BBB penetration of the gadobenate dimeglumine contrast (Fig. 2C and D), making this model more clinically relevant than those with leaky tumors. T2-weighted (T2w) MRI is the basic of all clinical diagnostic scans for DIPG patients and was employed in this study to determine the exact tumor location and volumes (Fig. 2C and D). The first control (no treatment) group showed a significant tumor progression (Fig. 2E). Our xenograft model showed a clinically relevant growth pattern marked by the ability to migrate through the cerebral spinal fluid and metastasize elsewhere in the central nervous system (Fig. 3E). H&E staining illustrated the focal region of growth with a diffuse tumor margin. After long periods of tumor growth, areas within the mass began to show regions of necrosis, most pronounced at the tumor center (pink arrow, Fig. 3F).

**Fig. 3.**
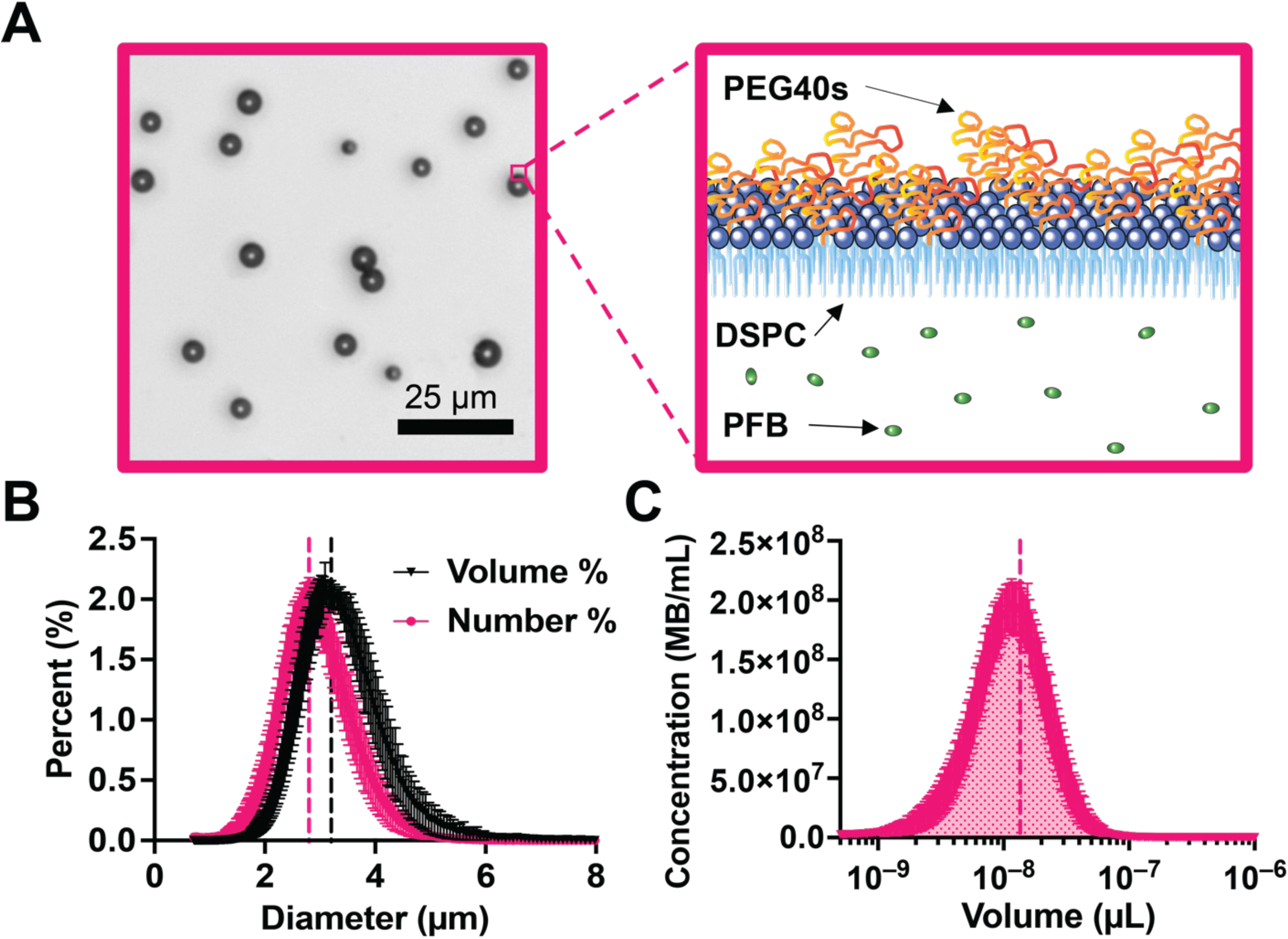
Microbubble characterization. (A) Brightfield image of 3 µm diameter microbubbles (left). Cartoon of microbubble structure showing lipid, PEG brush and internal gas (right). (B) Number- and volume-weighted size distributions. (C) Microbubble concentration against microbubble volume at a basis concentration of 10^10^ MBs/mL; shaded area under the curve represents the gas volume fraction. Vertical dotted lines represent mean values for both B and C. Data points represent the mean ± standard deviation (*n* = 8). Abbreviations: PEG = Polyethylene glycol; PEG40s = PEG_40_ stearate; DSPC = Distearoylphosphatidylcholine; PFB = Perfluorobutane; MB = Microbubble.

### Microbubbles are monodispersed in size and concentration

After the size isolation of 3 µm diameter microbubbles, the MB formulation was inspected using brightfield microscopy as shown in Fig. 3A. A Coulter Multisizer 4 was used to confirm concentrations and size distributions for all samples. The monodispersity of microbubbles was illustrated with narrow peaks for both number- and volume-weighted distributions (Fig. 3B). The resulting mean diameters were 2.8 and 3.2 µm, respectively. Each population was individually analyzed to confirm consistent microbubble volume dose injected^24^. Fig. 3C illustrates the microbubble concentration versus volume plot integrated to determine the gas volume fraction (𝜙𝑀𝐵). The mean 𝜙𝑀𝐵 for 10^10^ MBs/mL was 13.5 µL/mL (Table 1).

**Table 1.** Microbubble formulation characterization. Table shows the mean and mode diameter, standard deviation, d10 and d90 (where 10 or 90 percent of all MBs fall below this diameter), in the number-weighted graph (Fig. 4B). The mean diameter for the volume-weighted distribution (Fig. 4B) is also shown. The average gas volume fraction (𝜙𝑀𝐵) for 10^10^ MBs/mL is given in the rightmost column and found using Fig. 4C. All values were averaged over 24 separate measurements (8 samples).

| Size Distribution Characterization |  |  |  |  |  |  |
| --- | --- | --- | --- | --- | --- | --- |
| Number % Parameters | | | | | Volume % Mean | $\phi_{MB}$ @<br>$10^{10}$ MBs/mL |
| Mean | Mode | SD | d<sub>10</sub> | d<sub>90</sub> |  |  |
| 2.8 $\mu\text{m}$ | 2.9 $\mu\text{m}$ | 0.5 $\mu\text{m}$ | 2.2 $\mu\text{m}$ | 3.4 $\mu\text{m}$ | 3.2 $\mu\text{m}$ | 13.5 $\mu\text{L/mL}$ |

### BBB opening confirmation at the target site

Accurate and precise BBB opening at the target site was confirmed for our conditions by MRI. Figure 4A illustrates the contrast enhancement of gadobenate dimeglumine in the brain parenchyma owing to BBB disruption by FUS/MB. Similar to our drug, Panobinostat, this Gd contrast agent weakly binds to albumin and exhibits similar BBB penetration^42,46^. We saw no extravasation just prior to FUS/MB treatment, then a significant amount just after the application of FUS/MB. Images taken just prior to FUS/MB application the following week also showed no Gd extravasation, confirming closure of the BBB after the prior FUS/MB treatment. A subset of mice was injected with Evan’s blue dye to further confirm targeting and the extent of extravasation. Figure 4B illustrates dye extravasation (bottom) compared to the gadobenate dimeglumine extravasation (top), showing targeting and patterning to T1w CE images.

**Fig. 4.**
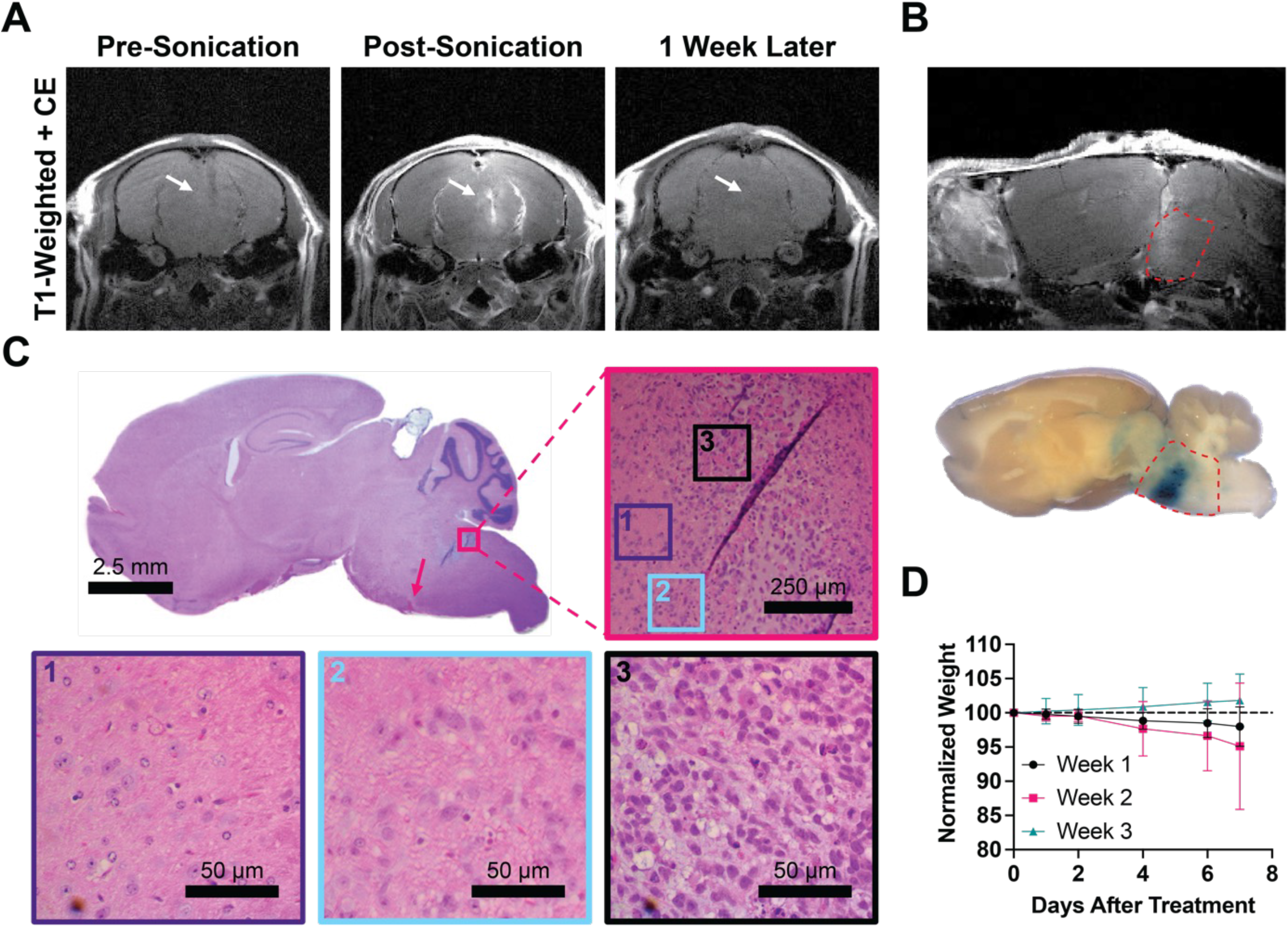
Confirmation of BBB Opening. (A) T1w CE MR images just before FUS/MB treatment (left), directly after treatment (center) showing BBB opening as bright spots owing to Gd extravasation and 1 week post FUS treatment (right) showing closing (white arrows); all images are in the axial plane. (B) Confirmation of BBB opening in the sagittal plane by T1w MRI (top), and Evan’s blue dye (bottom). Pontine region outlined in red dotted line. (C) H&E staining of the whole brain, with a pink arrow illustrating the region of red blood cell extravasation. Figure is zoomed (50x) into the region on the border of tumor (pink) and heathy tissue. Also shown are zoomed in regions of healthy tissue (purple border), tumor margin (blue border) and tumor (black border). (D) Plot showing the normalized body weight of each mouse after treatments. Data is split between weeks. There is no significant difference between each week.

Histological analysis was performed on a FUS/MB treated mouse to confirm the lack of morphological damage (Fig. 4C). H&E staining showed no major changes other than small amounts of red blood cell extravasation at the bottom of the FUS focal zone. After each treatment, mice remained within 94% of their initial body weight at days 1, 2, 4, 6 and 7 post FUS/MB treatment (Fig. 4D).

### Mild harmonic MB activity during FUS confirmed by PCD

During each FUS/MB treatment, a passive cavitation detector (PCD) recorded the acoustic response within the focal region. All voltage-vs-time signals were converted to the frequency domain for analysis (Fig. 5A). As expected, during treatments there was a strong harmonic response absent of significant broadband feedback, indicating mild MB harmonic oscillations without inertial implosions. During the initial time just after the injection of MBs, we saw a spike in harmonic cavitation dose (HCD), illustrated in Fig. 5B. As FUS application continued, there was a decay curve as microbubbles were cleared from circulation. Meanwhile, broadband cavitation dose (BCD) maintained a similar intensity before and after microbubble injection, indicative of general system noise but not MB implosions. Figure 5C shows the spectral content of a single 10 ms pulse with 0.5 ms readings before and after FUS. Beyond the fundamental frequency response, significant sub- and ultra-harmonics signals were observed only during treatment. These single pulses were connected (without 0.5 ms ends) to give an overall visualization of the treatment, showing that the sub- and ultra-harmonic components were maintained during the entire FUS treatment (Fig. 5D).

**Fig. 5.**
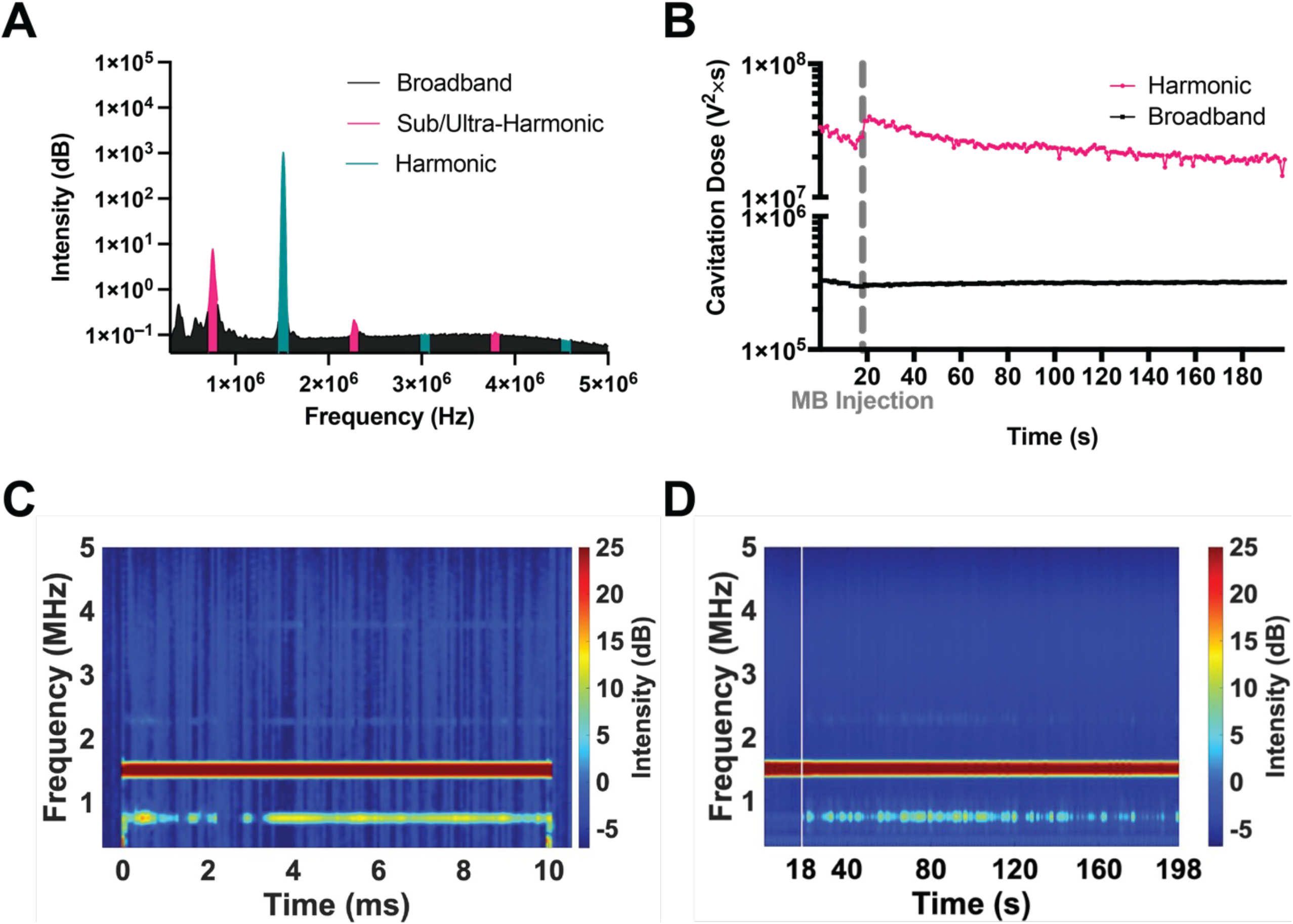
Passive cavitation detection of microbubble activity. (A) PCD voltage plot showing the area under the curve for sub/ultra-harmonics (pink), harmonics (teal), and broadband (black) peaks. (B) Cavitation doses calculated at each FUS pulse and plotted over the entire FUS treatment. Pink illustrates harmonic cavitation dose, while black shows broadband cavitation dose. A break in the ordinate was made to better view both curves. Each point represents the mean (*n* = 20). (C) Representative spectrogram of single 10 ms pulse with 0.5 ms recording before and after. (D) Representative spectrogram of a whole treatment where each 10 ms pulse is shown for each second (1 Hz pulse repetition frequency). Time in between FUS pulses is not shown. White vertical line represents the time when microbubbles were injected.

### MRI-guided FUS/MB improves tumor uptake of panobinostat

Once we confirmed BBB disruption was safe and effective at delivering gadobenate dimeglumine, we next confirmed targeted panobinostat delivery to the tumor. At 2 weeks post tumor injection, mice were treated with MRI-guided FUS/MB immediately followed by IP injection of panobinostat (10 mg/kg). Liquid chromatography with tandem mass spectrometry (LC-MS/MS) was used to determine the concentration of panobinostat in the tumor region and front part of the cortex. FUS/MB was associated with significantly more panobinostat in the tumor region (194.3 ng/g) than in the frontal cortex region (65.6 ng/g, Fig. 6A). Mice showed significantly more panobinostat in the tumor region than when treated with FUS/MB than without FUS/MB, with a mean concentration of 194.3 versus 61.8 ng/g (*p* < 0.0001, Fig. 6B). The tumor-to-serum ratio showed a significant increase between FUS/MB and non-FUS/MB mice, with a mean of 0.74 versus 0.18 mL/g (*p* < 0.05, Fig. 6C). Control mice treated with FUS/MB but without panobinostat showed no concentration of panobinostat in either the blood or brain tissue (Fig. S2).

**Fig. 6.**
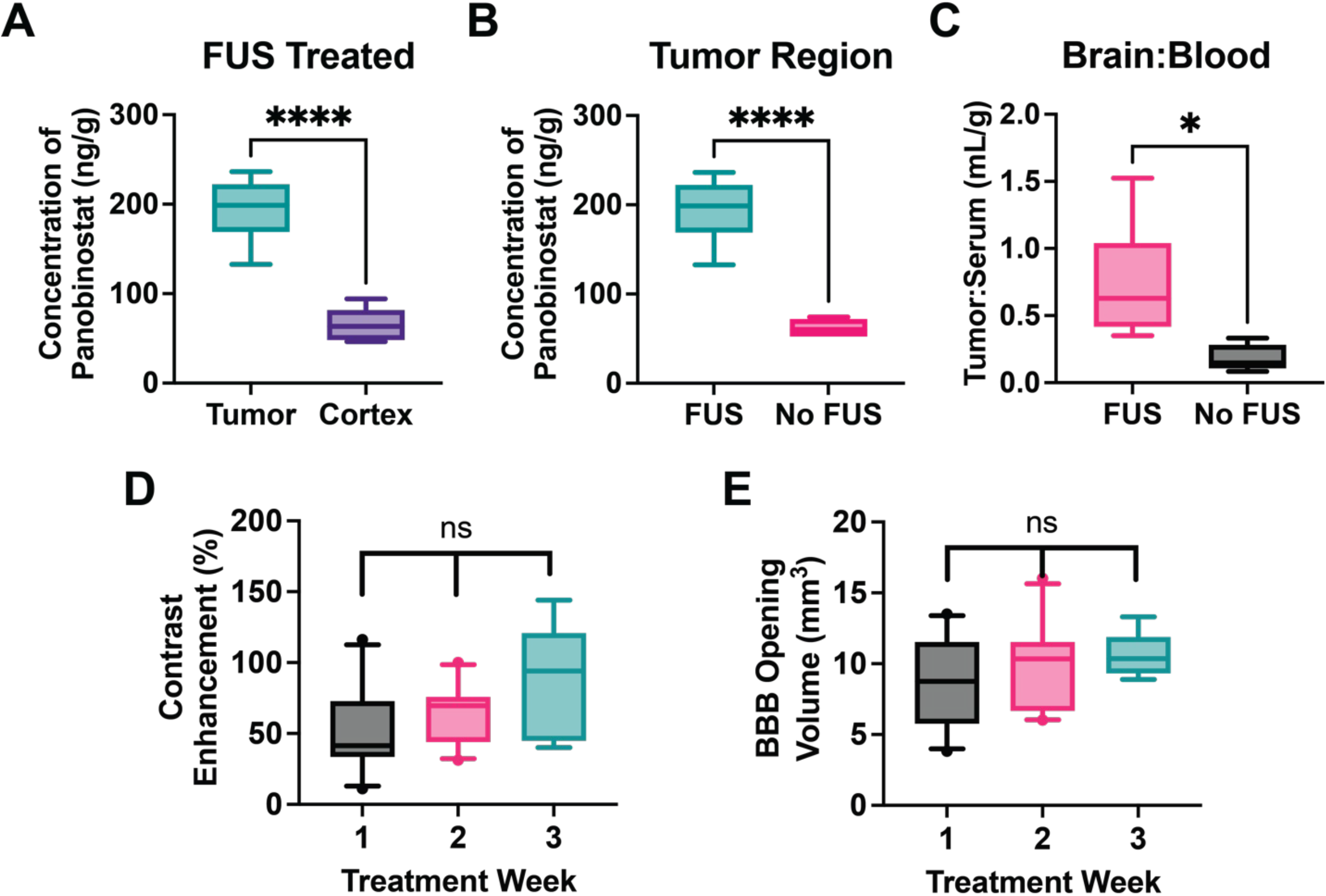
Increased delivery of panobinostat into DIPG tumors using MRI guided FUS/MB. (A) LC-MS/MS results showing panobinostat concentration after MB+FUS treatment in the tumor region vs. cortex. (B) Concentration difference between both tumor regions of FUS/MB-treated and untreated mice. (C) The tumor-to-serum ratio of panobinostat concentration for FUS/MB versus MBs alone (no FUS). (D) Contrast enhancement within the treated tumor region compared to the contralateral side. (E)Volume of BBB disruption determined by MRI T1w CE. Data represents mean ± standard deviation (*n* = 6 for A-C; *n* = 25 for D and E). Symbols *, **** represent *p* < 0.05 and *p* < 0.0001 respectively.

Prior to all treatments, T1w CE MR images were taken and analyzed for the extent of BBB opening volume and total contrast enhancement. The resulting BBB opening volumes were found to be 8.74, 9.68 and 10.56 mm^3^ as the weeks progressed (1, 2 and 3 respectively). Contrast enhancements for each were 51.5, 63.1, and 85 percent, respectively. No significant difference was observed in either volume or contrast enhancement during treatments week to week (Fig. 6D and 6E).

### FUS/MB and panobinostat is effective at reducing tumor growth and improving survival

Our data confirmed increased panobinostat delivery and its tumor cell cytotoxic effectiveness against DIPG in our murine model. During each treatment, three dimensional (3D)-T2w MR images in axial, coronal and sagittal planes were acquired to monitor tumor volumes (Fig. 7D). MR images were analyzed to find total tumor volume and monitor week to week (Fig. 7C) and showed that FUS/MB and panobinostat treatment inhibited tumor growth. MRI also showed significant intracranial edema, ventricle infiltration and inflammation in later time points. Tumor progression was significantly different between treated and untreated groups by week 3, where mean volume was 49.7 mm^3^ for non-FUS/MB and 14.1 mm^3^ for FUS/MB treated mice (*p* < 0.01). Neither group showed a significant drop in weight until week 3, when non-FUS/MB mice dropped to 88 percent of the starting weight. FUS/MB treated mice weight dropped to 91 percent of the starting weight, at most (Fig. 7A). Kaplan-Meier survival curves showed an increase in survival for the FUS/MB and panobinostat group with a mean survival of 31.5 days. This compared to the non-FUS/MB with panobinostat control group, which had a mean survival of 21 days. Immunohistochemical analysis of both treatment arms (panobinostat alone and panobinostat with FUS/MB) was conducted after a single treatment. Results showed that Ki-67 positive cells had decreased significantly in the FUS/MB treated group (Fig. 7E). Representative images are shown on the left side of Fig. 7E, and quantitative analysis is shown on the plot on the right.

**Fig. 7.**
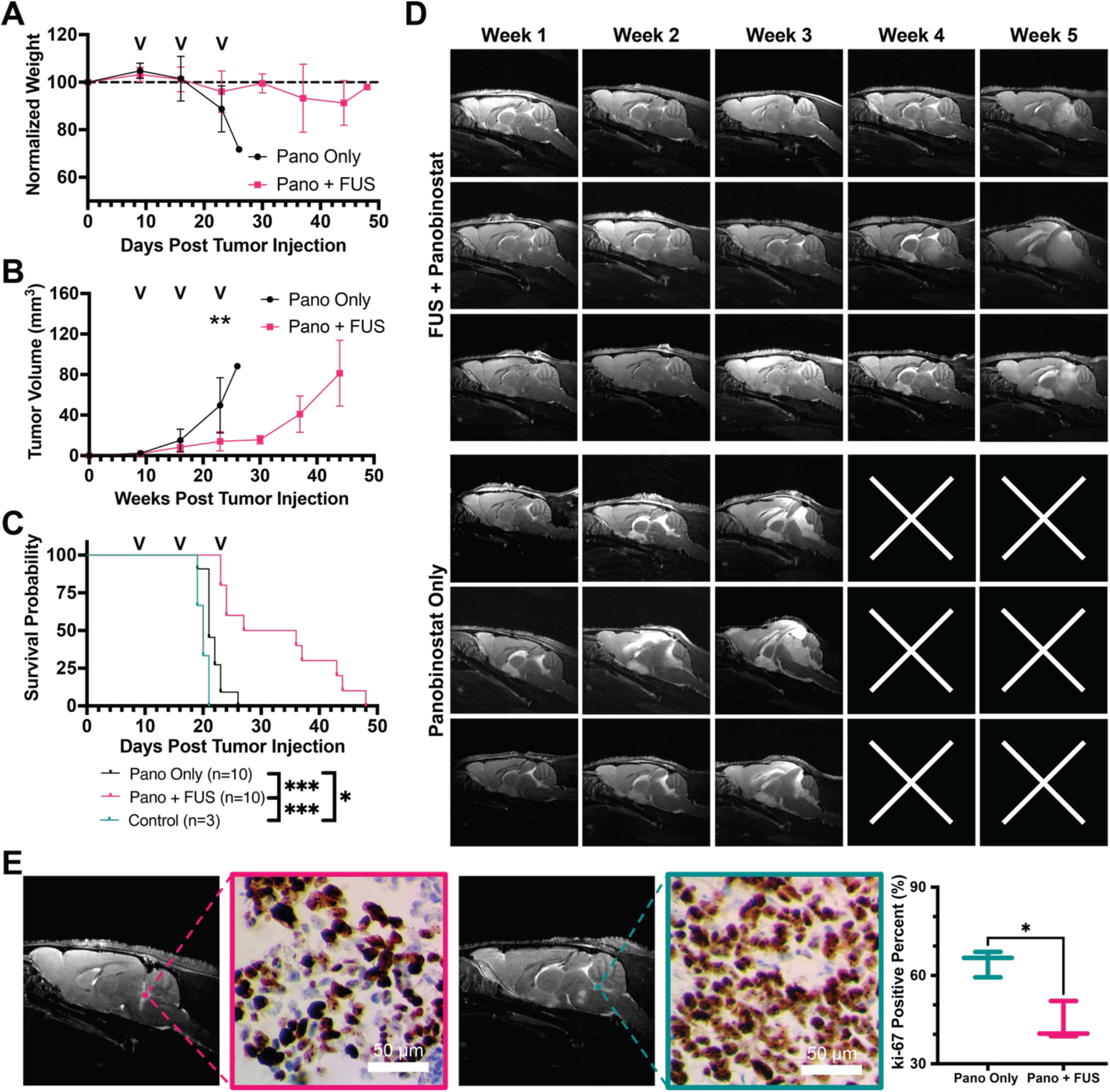
FUS/MB with panobinostat reduces tumor growth and improves survival. (A) Longitudinal study of mouse body weight obtained weekly, (B) tumor volume using T2w MRI, and (C) KP survival curves. Black arrows on plots represent the treatment days. (D) Representative T2w MRI (sagittal axis) images of tumor progression for 3 representative mice in each group. Top three rows show MB+FUS with panobinostat treated mice, and bottom rows are only panobinostat (no MB+FUS). White “x” illustrates the death of the mouse prior to the MR imaging that week. (E) Representative images of Ki-67 staining and its relation to MR images (left) and quantification of Ki-67 positive cells in tumor region for panobinostat + MB+FUS and panobinostat only (right). Significance testing was done using an unpaired Student’s t-test for B (*n* = 10) and E (*n* = 3), and Mantel-Cox test for C (*n* = 10), where *, **, *** indicate P <0.05, 0.01, and 0.001 respectively.

## DISCUSSION

While outcomes have improved for almost all childhood cancers, DIPG has shown little to no improvement in survival over decades. The tumor’s diffuse nature and location causes it to be unresectable and limits any ablation options^1^. Given the retention of the blood brain barrier (BBB) in DIPG, drug delivery has another hurdle^47^. This is evident on MRI, in which contrast enhanced images show little to no enhancement^2^. Many therapeutics have shown effectiveness *in vitro*; they just cannot pass the BBB in effective enough doses to treat the cancer without causing systemic toxicity^47^.

MRI-guided FUS with microbubbles (FUS/MB) has become a non-invasive way to temporarily disrupt the BBB, providing a window for therapeutics to enter only at the tumor site. FUS transducers can maintain a focal region on a millimeter scale and therefore can hit accurately and precisely only where the tumor is located. Several groups have looked into this technology to deliver drugs to solid tumors. Some of these trials have shown increased survival and reduced tumor growth^23,35,48–57^. As a consequence, this technology has been translated into clinical trials showing the safety of FUS/MB treatments, and current clinical trials are focused on efficacy with tumor types other than DIPG, including glioblastomas. Due to the depth and targeting complexity of the brainstem, only a few groups have investigated using this technology for pontine DMG^54,55,57–59^. Over the last 5 years, two groups have looked into using MRI-guided FUS/MB to treat DMGs, and one group used bioluminescence-guided FUS in pre-clinical models. Initially in 2018, Alli *et al.* showed the ability to deliver doxorubicin to the pons without any neurological issues. Later, Englander *et al.* used etoposide and treated for two consecutive weeks, confirming the safety of multiple treatments. To our knowledge, no group has yet shown a significant increase in survival for DMGs^54,57,58^.

In our study, we used a H327-altered, TP53-mutant DIPG PDX murine model to determine the efficacy and safety of MRI-guided FUS/MB treatments. We tested the HDAC inhibitor, panobinostat, due to previous drug screening studies using patient-derived DIPG cell lines^9^. Panobinostat specifically targets H3K27M cell populations, showing antitumor efficacy both *in vitro* and in xenograft models^60^. Overall, these studies showed impaired cell proliferation and viability of DIPG cells^9^. In cell lines with the H3K27M alteration, panobinostat reduced tumor growth and increased H3K27 trimethylation. While Hennika *et al.* saw epigenetic effects, unfortunately the systemic doses (20 mg/kg) resulted in significant toxicity. When the dose was reduced (10 mg/kg), it was well tolerated but failed to prolong survival^61^. Unlike our model, many of these prior models showed a slightly leaky BBB^61^. The molecular weight of panobinostat is relatively small (439.51 Da for the base form of panobinostat and 349.43 Da for the anhydrous lactate form)^42^, but most (∼90%) of the drug is protein bound in blood, which prevents its penetration across the BBB or BBTB^41^. As we showed, our cell line yields an intact BBB, which is more clinically relevant. Our objective was to use MRI-guided FUS/MB to obtain high enough doses at the tumor site while minimizing toxic side effects by injecting the lower dose (10 mg/kg). Using MR-guided FUS/MB, we demonstrated a significant increase in panobinostat concentration in the parenchyma at the tumor location compared to the cortex (3-fold), illustrating both the delivery enhancement, and targeting capability of the technology. We also found a significant increase in panobinostat in FUS/MB-exposed tumors over non-FUS/MB ones (over 3-fold). With similar serum values between groups, this led to a 4-fold increase in tumor-to-serum ratio at 1 hour post IP panobinostat injection.

Our study is the first to demonstrate three repeated treatments to the brainstem. Similar to Englander *et al.,* we saw reopening with repeated treatments^54^. We consistently had a BBB opening volume of ∼10 mm^3^ over all three weeks, which is just above the expected focal volume of the transducer (∼8.4 mm^3^). The volumes did not change significantly over the three weeks of treatment. Many research groups have begun to use PCD feedback to ensure a mild and consistent cavitation dose^34,35,62^. Our study used a constant MI (0.4) with a bolus injection of monodisperse microbubbles. Our PCD recording prior to microbubble injection showed the spike in harmonic oscillations as microbubbles entered circulation, and the harmonic cavitation dose decreased with subsequent clearance following the expected decay from prior pharmacokinetic analysis of similarly formulated and sized microbubbles^63^. The use of a PCD feedback treatment plan necessitates a continuous infusion of MBs, and hence was considered too time consuming for our purposes. The harmonic acoustic response is known to be associated with mild MB oscillations^33^, which is also known to be effective for BBB opening. We also avoided the broadband (inertial) acoustic response associated with MB implosions and unwanted biological effects^33,64^.

Despite encouraging results of the FUS/MB technique in glioblastomas^23,51,56^, a survival benefit with FUS/MB had not yet been shown in animal models of DMG. Here, we demonstrate the first significant survival benefit in a DIPG model using FUS/MB. We also show that FUS/MB-mediated BBB opening is accurate, precise, and reproducible in weekly treatments. Normalized body weight showed relatively stable weight for the FUS/MB treated mice. During the initial 4 weeks of the study, we found significantly decreased tumor growth in the FUS/MB-treated mice, showing the emergent potency of an otherwise ineffective anticancer drug by use of a noninvasive image-guided drug delivery technology. It is likely that the reduced tumor growth we observed was responsible for the survival benefit, in which we observed an increase in the average overall survival from 21 to 31.5 days for the FUS/MB-treated mice versus drug alone.

Our murine PDX model shows similar pathology, mutations, MRI features and growth patterns to DIPG in human patients. The MRI-guided FUS/MB technology was shown here to be a safe and effective way to provide a higher concentration of panobinostat at the tumor region without increasing systemic dose or off target effects. However, our study had some limitations. Our study was designed to confirm the safety and effectiveness of FUS/MB for only 4 weeks.

Although this is longer than other groups have studied, future work should investigate whether a longer-term survival benefit can be shown. Specifically, it must be investigated whether or not these DMG tumors can develop a resistance to panobinostat and/or FUS/MB over time. Other research with panobinostat would indicate that the former may occur^65^. Therefore, more research is necessary with other drugs and drug combinations in conjunction with FUS/MB. Additionally, our FUS transducer had a relatively small focal area, although we were still able to cover most of the mouse tumor region with a single pulse (>75%). Given the small size of the mouse pons, we observed this focal region overlap with healthy tissue during FUS/MB treatment. Given the scale change moving to a pediatric human patient, this would not be the case; a larger focal spot with multiple targets would be necessary and would be expected to target the tumor accurately if guided by MRI, e.g., by identifying the treatment margins with T2w MRI. Our center frequency of 1.515 MHz would have to be reduced as we would no longer need the tight focal region but instead require more penetration depth through a larger brain and the thicker skull. Finally, our exact drug formulation (free panobinostat) would have to change as it is no longer on the market. We would move to the water-soluble formulation (MTX-110) currently being used in conjunction with CED. Moreover, our work here is generalizable to other key drugs.

In conclusion, our group has shown that MR guided FUS/MB is a non-invasive, safe, and effective way to temporarily disrupt the BBB and increase drug delivery of panobinostat in a clinically relevant DIPG orthotopic xenograft mouse model. We showed the potential to treat this model weekly for 3 weeks while maintaining safe and similar BBB opening volumes. This is a critical pre-clinical step to inform the use of this technology for DIPG patients in human clinical trials.

## MATERIALS AND METHODS

### Experimental Design

Sample size for our survival study was calculated using a log rank test of power. Using an expected hazard rate of 0.17 we determined a sample size of n = 10 would give us 80% power. For the survival study, body weight was measured up to 5 times a week and mice were monitored daily and euthanized at endpoints which included irreversible neurological deficit, body condition score less than 2, or weight loss of greater than 20%. At this point mice were euthanized via CO_2_ inhalation. Mouse survival was closely monitored till the end of the experiments. Each animal was counted as a single replicate throughout the study.

Our study was conducted to determine if FUS+MBs could improve the delivery and efficacy of panobinostat in a DMG model. This was done within a controlled laboratory setting using mice. All experiments involving animals were conducted according to the regulations and policies of the Institutional Animal Care and Use Committee (IACUC). When selecting groups all mice were randomly selected with an even amount used from each cohort of mice deliveries.

### Cell lines and cell cultures

DIPG cell line, BT-245 (H3.3K27M-mutant pediatric DMG from Dr. Keith Ligon, Dana-Farber Cancer Institute), was biopsied from a patient at Boston’s Children Hospital. Cells were cultured on plates coated with poly-L-ornithine (0.01%) (Sigma) and laminin (0.01 mg/mL) (Sigma). Cells were grown in Neurocult NS-A media (Stemcell Technologies) supplemented with penicillin-streptomycin (1:100), heparin (2 μg/mL), human epidermal growth factor (EGF; 20 ng/mL), and human basic fibroblast growth factor (FGFb; 10 ng/mL) was used to maintain all lines.

### Cell Viability Assay

Cell proliferation and viability was examined using the MTS [3-(4, 5-dimethylthiazol-2-yl)-5-(3-carboxymethoxyphenyl)-2-(4-sulfophenyl)-2H-tetrazolium] assay. CellTiter 96 AQueous One Solution (Promega, Madison, WI, USA) were used. Cells were seeded at 20,000 cells per well into a 96-well plate (Corning) in neurosphere culture, in a media volume of 100 µl. Twenty-four hours later, the cells were treated with a range of doses of panobinostat (Selleckchem, Houston, Texas, USA) in triplicate. At the end of the drug treatment period (72 h), 20 µl of MTS reagent was added to each well to make a final volume of 120 µl. Absorbance values for plate wells were determined using a BioTek Synergy 2 plate reader at a wavelength of 490 nm after 3 h of incubation. For all tests the background absorbance was subtracted. IC50 values were determined experimentally through Prism 9.

### Microbubble Preparation

Lipid-coated MBs with a Perfluorobutane (PFB) gas core were prepared using sonication, as described previously by Fesitan et al.^66^. Under completely sterile conditions, a lipid film comprising DSPC: PEG40S (90:10) was hydrated with filtered and sterile PBS (1X) to a final concentration of 2 mg/mL at 65 °C for 40 min. Once fully hydrated, the solution was cooled to room temperature. The lipid solution was sonicated with a 20 kHz probe (model 250A, Branson Ultrasonics; CT, USA) at low power (3/10; 3 W) for four minutes. Again, allowed to cool to room temperature, PFB was delivered to the surface of the lipid suspension for 15 s. The solution was then sonicated at high power (10/10; 33 W) for 30 s to yield polydisperse MBs. Polydisperse MBs were then collected into 30 mL sterile syringes and isolated by differential centrifugation into 3 ± 0.5 µm in diameter. The size isolation process, including time and speeds of centrifugation, can be found in Fig. S3. MB concentration and number- and volume-weighted size distributions were measured using a Multisizer 3 Coulter Counter (Beckman Coulter). MB concentration (𝑐_i_, MBs/µL) versus MB volume (𝑣_i_, µL/MB) was plotted, and MB gas volume fraction (𝜙𝑀𝐵) was estimated as follows:

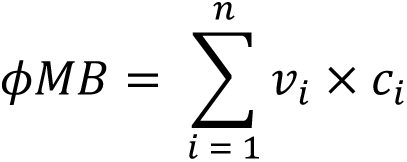

where 𝑖 is the index of the sizing bin, and there were 300 bins ranging from 0.7 to 18 µm in diameter. Three independent MB preparations were measured after synthesis and two hours prior to FUS treatment to confirm size distributions and concentration. MB cakes were stored in the refrigerator at 4 °C for use at a later time.

### Orthotopic PDX mouse model

Female athymic nude (nu/nu) mice (Charles River Laboratories, Wilmington, MA, USA) between 6 and 8 weeks old were initially anesthetized under isoflurane (3.5 %) then transferred to stereotactic set up and continued on isoflurane (1.8 %). A subcutaneous injection of carprofen (5 mg/kg) was given on the right flank. Head was sterilized with a betadine (5 %, Purdue, Stamford, CT, USA) wash followed by an ethanol (70 %) wash three times. A small incision (2 mm) and burr hole (1 mm in diameter) was created on the top of the skull at 1 mm right and 0.8 mm posterior to Lambda landmark. A suspension of ∼200,000 BT245-luc2-GFP (H3.3K27M mutant) cells in a total volume of 3 μL serum-free media was injected at a rate of 600 nL/min into the brain 5 mm deep from the skull at the craniotomy location. After injection, bone wax (SMI, Saint-Vith Belgium) was used to fill craniotomy and sutures were used to close incision. For pain management, a topical analgesic and antibiotic was rubbed on the incision site. Mice were given carprofen (5 mg/kg) at 24 and 48 hours post injection. Each mouse was closely monitored for any recovery issues. Mice were randomly assigned to each treatment group.

### Magnetic Resonance Imaging

A Bruker Biospec 9.4 Tesla MR Scanner (Bruker, Billerica, MA) with a mouse head phase array coil was used for all MR images. Prior to MRI, mice were anesthetized with 1.8% isoflurane and monitored throughout the imaging. Mice were placed into modified MRI bed that prevented movement of mouse head during transfer from MRI to focused ultrasound (FUS) system and inserted into the MR scanner. First, a localizer scan was acquired brain localization. For the FUS treatment group, a T1w fast spin echo RARE (*Rapid Acquisition with Relaxation Enhancement*) images were acquired in the axial plane (repetition time (TR)/echo time (TE), 720/12 ms; flip angle, 90°; number of averages, 4; field of view, 20 mm × 20 mm; resolution, 78 μm x 78 μm x 700 μm) was performed 12 min after intravenous injection of 0.2 mmol/kg gadobenate dimeglumine (MultiHance, Bracco, Princeton, NJ). All mice underwent a high-resolution 3D T2-turboRARE scans in sagittal, coronal, and axial planes (repetition time (TR)/echo time (TE), 2511/33 ms; flip angle, 90°; number of averages, 4; field of view, 20 mm × 20 mm; resolution, 78 μm x 78 μm x 700 μm). Mice remained stereo tactically placed on MRI bed and transferred to FUS system for treatment where an intravenous injection of 0.1 mL gadobenate dimeglumine (MultiHance, Bracco, Milan, Italy) was given. After treatment, mice were moved back to MRI and a post FUS T1-weighted sequence (same parameters as previous T1) was acquired after 12 minutes post gadobenate dimeglumine injection. All image acquisition was performed using Bruker ParaVision NEO360 v.3.3 software.

### MR Image Analysis

T1-weighted images were used to quantify extent of blood brain barrier disruption using FIJI (Maryland, USA). First a region of interest (ROI) was defined within contralateral side of the brain (left) in order to determine the baseline intensity. The area of BBBD was defined by the control region plus 2 standard deviations. The area found was calculated on all MRI slices and volume was totaled by multiplying the slice thickness by the average area of adjacent slices and summed for all slices. The contrast enhancement was determined by determining the average intensity within BBBD volume and dividing by the intensity of the control region. T2-weighted MR images were used to calculate tumor volumes. Three axis images were uploaded to Slicer^67^. Tumor margins were defined on each slice then a total volume was calculated by summing each slice volume and multiplying by slice thickness (0.7 mm). This was completed on both sagittal, and axial slices and the average volume was taken.

### MRIgFUS and Drug Treatment

BT245 xenografted mice were allowed to grow tumors for 9 days. At this point weekly treatments of panobinostat (10 mg/kg) began and continued for 3 weeks. The experimental set up is shown in Fig. 1B. A single element, geometrically focused transducer (center frequency = 1.515 MHz, diameter = 30 mm) was driven by the RK-50 system (FUS Instruments, Toronto, Canada). Concentrically within the FUS transducer a single element, geometrically focused transducer (center frequency = 0.7575 MHz, diameter = 10 mm) was used for passive cavitation detection where voltage data was collected on the RK-50 system. Using the pre-FUS T2-weighted MR image (coronal axis), the tumor center was targeted (Fig. 1.1). Ultrasound gel was placed on the mouse head and lightly scrapped in to remove any air bubbles between the skin and gel. An acoustically transparent tank filled with degassed waster was placed on top of the gel ensuring no air bubbles were trapped Fig. 1B. Microbubbles (25 μL/kg; diluted in 0.1 mL) and 0.1 mL gadobenate dimeglumine were injected intravenously through a tail vein injection via 26 Ga needle. Just prior (10-20 seconds) to injection FUS was applied to determine baseline acoustic response without the presence of microbubbles. FUS parameters were as follows: 10 ms burst length, 1 Hz pulse repetition frequency, 3-minute post injection FUS treatment time, and a peak negative pressure of 0.615 MPa. Voltage data from the PCD was collected during the entire FUS treatment and analyzed as previously described^32^. Remining PCD analysis was done using MATLAB (Massachusetts, USA) including the calculations of harmonic and broadband cavitation doses. Directly after FUS treatment, panobinostat (10 mg/kg; diluted in 0.125 mL solution of 4% DMSO, 46% PEG300, and 50% PBS) was injected intraperitonially (IP). Mice where then sent back to MRI to complete post FUS T1 imaging.

### Liquid Chromatography-tandem Mass Spectrometry (LC-MS/MS)

A subset of mice (*n* = 18) was kept for 16 days post BT245 cell injections to allow extensive tumor growth. One third of the mice were treated with MRIgFUS and panobinostat IP injection, another one third were imaged on MRI and injected with panobinostat IP, the final group was untreated. At 60 minutes post panobinostat administration, mice were anesthetized under 4% isoflurane and blood samples were collected via cardiac puncture. Immediately after blood collection, a transcardial perfusion with PBS (1X) was conducted for 3 minutes. The brain was extracted and dissected axially at the midline between the cerebellum (pons and tumor area) and forebrain (cortex). Brain tissues were immediately snap-frozen using liquid nitrogen. Blood was allowed to clot for 45 minutes then centrifuged at 2000xg for 15 minutes. Serum (supernatant) was collected and snap-frozen using liquid nitrogen. All tissues were stored at -80 °C until LC-MS/MS analysis. Samples were analyzed by the School of Pharmacy and Pharmaceutical Sciences at the Anschutz Medical Campus. The analysis was performed on an Applied Biosystems Sciex 4000 (Applied Biosystems; Foster City, CA, USA) equipped with a Shimadzu HPLC (Shimadzu Scientific Instruments, Inc.; Columbia, MD, USA) and Shimadzu auto-sampler. An extend-C18 Zorbax column (Agilent Technologies) 4.6 x 50 mm, with a column guard held at 40°C was utilized. Solvent A: HPLC H2O with 10 mM NH4OAc and 0.1% formic acid; and solvent B: 1:1 methanol: acetonitrile. In triplicate, a 16-point standard curve of panobinostat was prepared starting from 20.0 µM and serially diluted. Samples were transferred to a 96-well plate and analyzed by LC/MS-MS method; 40 µL sample sizes were injected onto the column. The LOD was 210 pg/mL and the LOQ was 850 pg/mL.

### Immunohistochemistry and Histology

A subset of mice (*n* = 7) underwent histological analysis. Tumors were allowed to grow for 2 weeks and treated with FUS + panobinostat (*n* = 3), or panobinostat only (*n* = 3). A final mouse was analyzed after 3 weeks of tumor growth without any treatments. Six hours after treatment (if applicable) mice were sacrificed and perfused with 10% formalin (Thermo Fisher Scientific) for 3 minutes. Brains were immediately dissected and put into 10% formalin solution overnight.

Tissues were processed and paraffinized in FFPE blocks and sectioned at a thickness of 5 µm. Next, samples were deparaffinized and antigen retrieval was performed in citrate buffer (pH = 6.0) for 15 minutes. Blocking was done with 10% goat serum and incubated for 1 hour at room temperature. Primary antibody staining was done with anti-Ki-67 1:100 (ab15580, Abcam, Cambridge, United Kingdom) and samples were incubated overnight at 4 °C. After 3 washes, a secondary antibody (anti-rabbit IgG, 1:1000, Invitrogen) was used and incubated for 1-hour at room temperature. After 3 more washes, signals were detected using DAB stain and counterstained with hematoxylin. Slides were mounted with Permount (Thermo Fisher Scientific). Microscope slides were imaged on a brightfield microscope at 1x, 10x, 20x, and 50x. Quantification of images was done in FIJI (NIH, Maryland, USA) where a color deconvolution was used for DAB and hematoxylin staining. Each deconvolution image was converted to binary where a threshold was determined to differentiate all cells in the image. The Analyze Particles function was used to count cells. The percentage of positive nuclei was done by dividing the stained cells by the total cell count. Each slice was analyzed in 3 separate fields of view within the tumor region.

### Statistical Analysis

All data collected is presented as mean ± standard deviation. No preprocessing was done to data with the exception of voltage data collected from the PCD. PCD data was preprocessed as described in Martinez et al^32^. All statistical analysis was completed in Prism 9 (GraphPad, California, USA). Star representations of p-values are indicated in captions and less than 0.05 was indicative of statistical significance. An unpaired Student’s t-test and Kaplan-Meier estimates were used to compare two groups and survival analysis respectively.

## Acknowledgments

The authors would like to thank John Desisto for his expertise throughout this project. Parts of figures 1 and 2 were made using Biorender.com.

## Author contributions

Conceptualization: NS, AG, MB

Methodology: PM, GN, JS, MFW, AP, JaS, KHS, NE, NS, AG

Investigation: PM, GN, JS, AP, BB, MS, AM, JaS, NS Visualization: PM, GN, JS, JaS, NS

Funding acquisition: NS, AG, MB Project administration: NS, AG, MB Supervision: NS, AG, MB

Writing – original draft: PM

Writing – review & editing: NE, NS, AG, MB

Competing interests: The authors declare that they have no competing interests.

Data and materials availability: All data are available in the main text or the supplementary materials.

