## Supplemental Materials for "MRI-Guided Focused Ultrasound Blood-Brain Barrier Opening Increases Drug Delivery and Efficacy in a Diffuse Midline Glioma Mouse Model"

**List of Supplementary Materials**

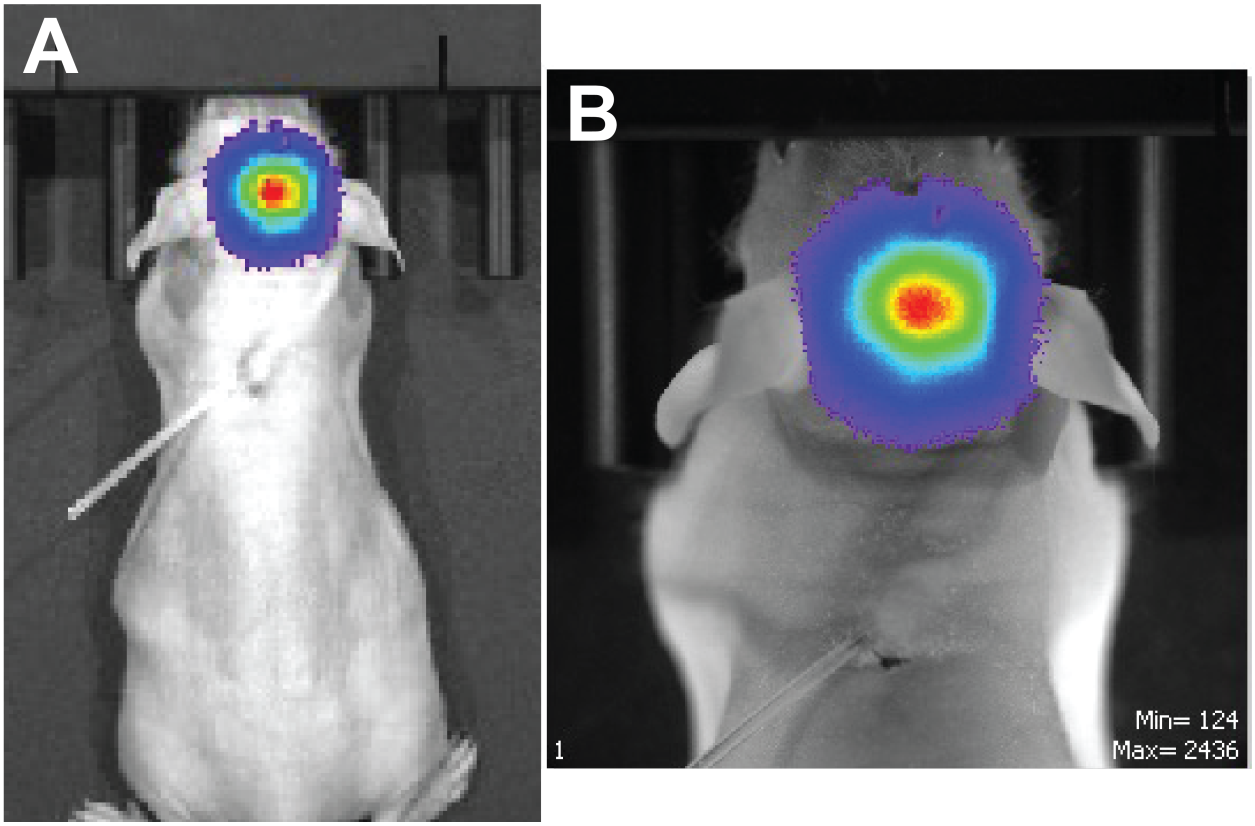


**Fig. S1: IVIS imaging of BT245 orthotopic xenografts.** Mice were injected with luciferin (150 mg/kg) IP and imaged on IVIS Spectrum. Full body (A) and head (B) scan with tumor indicated by bioluminescence intensity.

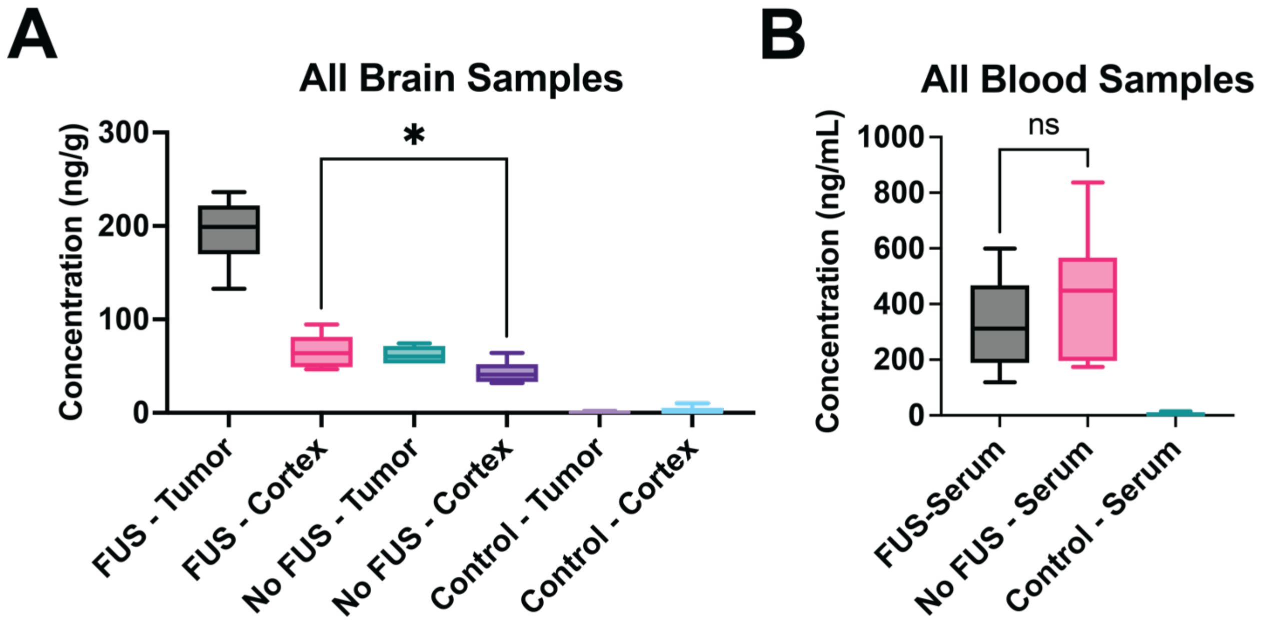


**Fig. S2: Liquid chromatography tandem mass spectrometry / mass spectrometry full results.** LC-MS/MS results of panobinostat concentration in FUS treated mouse brain (A) and blood (B). Data is represented as mean ± standard deviation ($n = 6$). Symbol * represents $p < 0.05$.

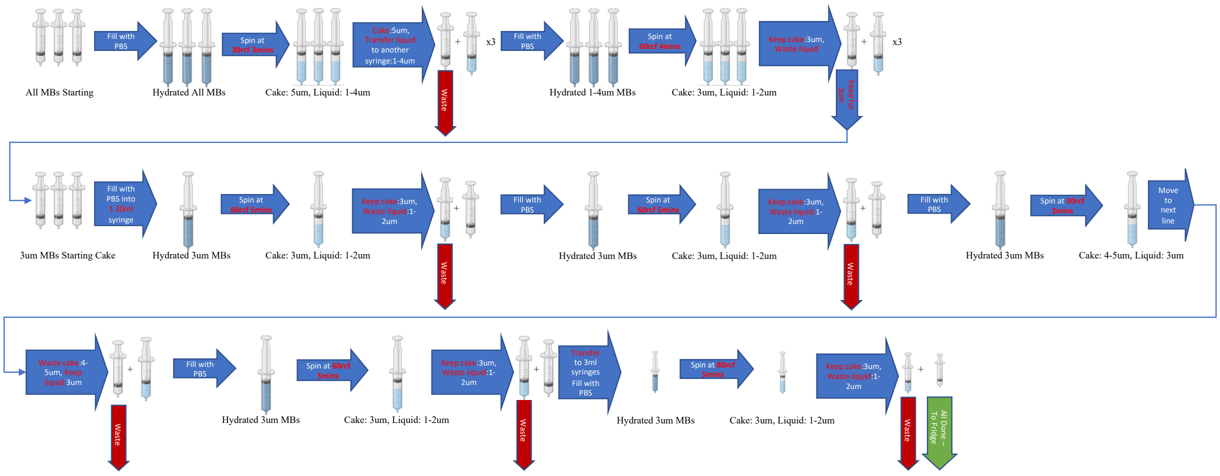


**Fig. S3: Size Isolation Process of Microbubbles.** This is an illustration of the process used to isolate the 3 µm size distributions used in the study. Thicker blue arrow illustrates separation of phases or centrifugation. Green arrows show when cake was to be saved for measuring.
